# Surveillance and molecular characterization of banana viruses and their association with *Musa* germplasm in Malawi

**DOI:** 10.1101/2024.06.24.600487

**Authors:** Johnny Isaac Gregorio Masangwa, Nuria Fontdevila Pareta, Philemon Moses, Eva Hřibová, Jaroslav Doležel, Isaac Fandika, Sebastien Massart

## Abstract

Malawi has local banana germplasm preferred by its population. However, the epidemics of banana bunchy top disease, caused by the banana bunchy top virus (BBTV), is wiping out the most preferred germplasm and limiting its plantation. On the other hand, there needs to be more knowledge on viruses present in Malawi on banana crops. Therefore, a survey was conducted country-wide to characterize banana germplasm and evaluate the presence, incidence and prevalence of banana viruses. The survey covered four country zones, combined farmer’s structured questionnaire and plant sampling. PCR products from infected germplasm were sequenced and aligned for each detected virus to build a phylogenetic tree. BBTV, BanMMV and BSV species were detected in Malawi. Malawi’s BBTV isolates belonged to the Pacific Indian Ocean group only and BanMMV isolates clustered to three sub-branches. All the BSV species present in Malawi belonged to clade 1. Among the genetic groups of Musa, the identified banana germplasm belonged to AA, AAA, AB, AAB, and ABB groups with some germplasms unique compared to already genotyped germplasms. The ABB group was dominant in Malawi and was often infected by BSV species (probably originating from endogenous viral sequences), while BBTV more often infected the AAA group. Banana propagule sharing was the primary source of banana planting materials with a higher risk of spreading virus diseases. The survey underlined the need to set up a banana seed industry and policies that promote farmers’ access to virus-tested planting materials, which will consequently prevent future virus epidemics.

## Introduction

Banana is an herbaceous plant belonging to the *Musa* genus and the *Musaceae* family [1]. It is ranked as the fourth most important food commodity in the world and represents a staple food and cash crop for millions of people in many developing countries [1, 2]. The crop possesses great genetic diversity in terms of its genomic groups based upon the inter- or intra-specific hybridization of *Musa acuminata* and *M. balbisiana*. The hybridization resulted in the development of AA, AB, AAA, AAB, ABB, AABB, AAAB, or ABBB genotypes [3]. Even though no African country is among the top five world producers of banana, countries such as Uganda, Rwanda and Cameroon have the highest per capita annual consumption, exceeding 200 kg and providing up to 25 percent of the daily calorie intake [4]. Banana is also an important food and cash crop in Malawi. Over 30% of Malawi’s population depends on banana cultivation for their livelihoods. In Malawi, like in many parts of the world, a large proportion of banana fruits are produced in small plots or backyard gardens [5, 6]. A study conducted in the early 2000s revealed that in Malawi’s banana major growing districts, 50% of farmers’ income came from banana [7] despite the presence of constraints such as drought, poor soil fertility, poor management, pests, and diseases [8].

Nevertheless, the situation dramatically changed by 2015, as 90 % banana plantations in the southern part of Malawi were wiped out by banana bunchy top disease (BBTD) caused by banana bunchy top virus (BBTV) [9]. BBTV is a member of genus *Babuvirus and* the family *Nanoviridae* whose isolates are divided into two major groups: the Pacific– Indian Oceans (PIO) group and South-East Asian (SEA) group. The latter group is confined in Asian region whilst the former is widely distributed[[10, [11]. According to Mikwamba *et al*. [12], Thyolo district, in Southern Malawi, was the worst hit district, with huge economic losses inflicted on many smallholder farmers’ livelihoods due to BBTD. Currently, BBTD is destroying banana plants also in Karonga, which is one of the major banana and plantain growing districts of northern Malawi [13], and is threatening the extinction of banana industry [14]. The Food and Agriculture Organization [15] estimated that BBTV alone has accounted for almost 40 % of loss of banana production in Malawi and Jekayinoluwa and colleagues [16] indicated that this disease has affected 80 % of banana producing areas in Malawi. This banana disease has not only affected farmers but also value chain players such as transporters, traders, agro-processers, researchers, micro-finance institutions and consumers [17]. BBTD caused the country to rely upon import of over 20,000 tons to meet the high demand [Ministry of Agriculture - unpublished]. The presence of BBTD also forced the country to source over 1,045,250 disease-free tissue culture planting materials from other countries [Ministry of Agriculture - unpublished] at an approximate cost of 2,086,500 US$.

Beyond BBTV, other viruses can cause problems in banana and other vegetatively propagated crops [18]. There are up to 20 different virus species reported to infect banana worldwide [1] causing different symptoms. For instance, banana streak viruses cause leaf chlorosis, necrotic streaks on leaves, stuntedness, fruit distortion, smaller bunches and splitting pseudostem, which leads to death of infected plants [19, 20, 21].

Banana streak viruses (BSV) belong to the genus *Badnavirus* in the family *Caulimoviridae*. There are nine BSVs species recognized by ICTV: *Banana streak OL virus* (BSOLV), *Banana streak MY virus,* (BSMYV), *Banana streak IM virus* (BSIMV), *Banana streak GF virus* (BSGFV), *Banana streak UA virus* (BSUAV), *Banana streak UI virus* (BSUIV), *Banana streak UM virus* (BSUMV), *Banana streak VN virus* (BSVNV), *Banana streak UL virus* (BSULV). The unclassified tentative species are *Banana streak CA virus* (BSCAV), *Banana streak UJ virus* (BSUJV), *Banana streak UK virus* (BSUKV), and *Banana streak PE virus* (BSPEV) [22]. Some BSV sequences do integrate full length of functional viral genome called infective endogenous sequences and some of the they do integrate in both most banana genotypes [23]. BSV can cause yield losses of 6 % to 15 % in banana crops depending on the variety of the banana, strain of the virus, number of the strains infecting the crop and the prevailing environmental factors [24, 25].

Banana mild mosaic virus (BanMMV) is a single stranded RNA (ssRNA) virus belonging to the *Betaflexiviridae* family, genus *Banmivirus.* This virus is the causative agent of Banana mild mosaic disease [26, 27] and is suspected to be spread mainly inadvertently in infected asymptomatic planting material, and no natural vector has been identified for BanMMV. It is nevertheless not widely recognized as a threatening pathogen of banana [20] but it contributes to symptom expression of necrotic streaks when present in a mixed infection with CMV [28].

*Cucumber mosaic virus* (CMV) has a very broad host range and is endemic to most of the banana growing regions [29]. Chlorosis, mosaic, heart rot stunted growth and low yields are symptoms associated with CMV [29, 30].

Banana bract mosaic virus (BBrMV) is identified by spindle shaped streaks, stripes on the pseudostem and mosaic pattern on the bracts. Distinct discontinuous streaks along the primary leaf veins which look irregularly thickened or raised are symptoms caused by a severe infection. Scattered white to yellowish streaks across from the midrib to the margin of the leaf are additional symptoms. The infected plant has an unusual reddish-brown or necrotic discontinuous streak towards the base of the pseudostem [31, 32].

Banana viral diseases are spread through vegetatively propagated plant materials [24, 33, 34] or either transmitted by vectors such as aphids (*Pentalonia nigronervosa* Coquerel and *Pentalonia caladii* van der Goot.) for BBTV [35], aphids (*Pentalonia nigronervosa)* for BBrMV [36], mealybugs (*Saccharicoccus sacchari* (Sugarcane mealybug), *Planococcus citri* (Citrus mealybug), Oleander mealybug and *Dysmicoccus brevipes* (pine apple mealybug) for BSVs [37; 38]. No natural vector has been identified for BanMMV.

So far, only BBTV has been officially reported in Malawi. No study has been done to investigate other banana viruses that are present in Malawi, their hotspot areas, and prevalence among Malawi’s banana cultivars. Lack of knowledge of other banana viruses may lead to a rapid distribution of banana planting materials infected with other viruses which could result in an outbreak of other viral diseases. With the foregoing in mind, the study was executed to evaluate the presence and incidence of banana viruses in *Musa* crops through a survey and targeted molecular detection of the viruses at country-wide level, and plot maps of all spots of banana viruses’ endemic areas. The relationship between genotypes, source and age of banana mat and virus infections were investigated. The surveyed area was divided into four regions that are referred to as clusters in this study.

## Materials and methods

### Banana viruses survey

A survey was conducted in August 2020 across Malawi in four banana cultivation zones. A total of 180 farmers were interviewed and 271 *Musa* leaf samples were collected. Zone 1 corresponded to the Southern region districts of Chikwawa, Mulanje, Nsanje and Thyolo; zone 2 included the Eastern region districts of Phalombe, Machinga, Mangochi and Zomba; zone 3 was the Centre region with Dedza, Lilongwe, Salima and Nkhotakota districts; and zone 4 was the North region of Chitipa, Karonga, Rumphi and Nkhata-bay districts. The listed sites laid within latitude -9.597858 S to -17.102709 S and longitude 33.219222 E to 35.814582 E. The areas’ altitude ranged from 48 to 1,665 meters above sea level (masl). Leaf samples were collected according to Kumar *et al*. [39]. Sampling sites were at a minimum of 5 km from each other. Collected leaves were dried over silica gel. Interviews and field observation allowed to gather data, that was recorded on the structured questionnaire that was uploaded on Mobile Data Collection (MDC.gis) software (https://mdc.giscloud.com).

### Targeted detection

40 mg of leaf samples were ground in 4 ml of CEB extraction buffer [40] in the Agdia extraction bags (Agdia-EMEA, Soisy-sur-Seine, France) using a tissue homogenizer (Agdia, Elkahart, USA). Depending on the species of the virus, immuno-capture PCR or immune-capture RT-PCR was done on the collected leaves. Tubes were coated with antibodies as describe by De Clerck *et al.* [41], then proceeded by adding 25 µl of plant extracts. The standardized protocols of the *Musa* Germplasm Health Unit (from the University of Liège) described by De Clerck et al. [41] were applied for the detection of viral species infecting banana. Complementary DNAs were prepared for RNA viruses. For DNA viruses (eg. BSV) DNase I (Invitrogen, stock 277 U/μl) treatment was done. Virus specific PCRs were prepared following the protocol followed by visualization of amplicons on agarose gel.

### Genotyping of Malawi’s banana cultivars

Local *Musa* landraces were collected from across Malawi for molecular characterization to understand its genetic diversity and conserve them in the International Genebank for future resilient and breeding programs. Genomic DNAs of banana germplasms were isolated from the cigar leaf and subjected to microsatellite (SSR) genotyping following Christelová *et al*., [42]. The SSR data of Malawi’s germplasms were analyzed together with the *Musa* core set collection [43] by DARwin software [44] and the cladogram was depicted with FigTree v1.4.4 (http://tree.bio.ed.ac.uk/software/figtree/). Ploidy level was determined using flow cytometry (FCM) according to Doležel *et al*. [45] and Christelová *et al.* [42].

### Map plotting and data analysis

Data collected during field survey and from the virus indexing were combined, then plotted using ArcMap in ArcGIS 10.5 software [46]. Chi-square test and analysis of variance were done to analyze survey data using STATA 16 Statistical package (https://www.stata.com). Chi-square test was used to analyze significant differences by sources of mat, banana genotypes, age of mats or cultivation systems. Analysis of variance was done to analyze the effects of sources of mat, banana genotypes, age of mats and cultivation systems depending on BBTV, BanMMV and/or BSV detection.

### Sequencing and phylogenic comparison of banana viruses present in Malawi

PCR products were purified using Qiaquick purification kit (Qiagen) according to the manufacturer’s instructions. The purified PCR products were then sequenced at Macrogen Europe using the specific primers (S1 Table). The sequences used for the phylogenic comparison were the partial DNA-R gene for BBTV, Coat Protein (CP) for BanMMV and ribonuclease H (RNAse H) for BSV from our study; and sequences that were downloaded from NCBI’s Nucleotide repository (S6 Table). The BBTV South Pacific group and the Asian group of isolates were used as references. BSV sequences from three BSV clades as published by Chabannes *et al.* [23] were used as references for BSV.

Different bootstrap maximum likelihood phylogenic trees replicated 1000 times were constructed using MEGA X software (https://www.megasoftware.net) to determine the homology and evolutionary relationships of the viruses in each group (BanMMV, BBTV, and BSV). Using the p-distance method the distances of the isolates were analyzed to understand their evolutionary history.

## Results

### Collection and molecular characterization of banana cultivars

During the survey, 273 leaf samples were collected from 17 districts of Malawi, corresponding to four banana cultivation zones. The farmer interview indicated that leaf samples originated from 38 banana cultivars. Genetic diversity of the Malawian genotypes conducted by SSR markers [42] was analyzed together with the banana core subset, characterized previously by Christelová et a. [43]. Banana core subset contained 591 germplasms, representing individual species/subspecies and subgroups of *Musa*, enabling a detailed genetic characterization of unknown banana genotypes [43, 47, 48]. The molecular characterization revealed that Malawi’s banana landrace germplasms belonged to 5 genomic groups, and all analyzed Malawi germplasms clustered together with the banana hybrids only (S2 Table). Twenty-two Malawian germplasms represented triploid AAA bananas, related to Cavendish, Rio, Red and Lujugira/Mutika groups of bananas.

Two Malawi germplasms (Mthwika and Suweshi) represented diploid AA Mchare bananas, one of the progenitors of triploid Cavendish and Gros Michel edible banana clones. Seven Malawi germplasms belonged to Plantain and Silk groups (AAB genome), and three Malawi germplasms represented Monthan and Pisang Awak banana groups with the ABB genome (Fig 1, S2 Table). Compared to the existing diversity, 4 germplasms were unique: Mabere (ABB) was distant from other ABB germplasms while Munowa; Kapeni and Ndifu germplasms were also distant from other AAA genotypes (Fig 1). The analysis also showed genotype ABB was the most sampled (59 %; n=160/271) (Table 1), whatever the zone (from 46% to 74%), followed by AAA at 25 % (n=68/271). Highly significant differences (P=0.000) were observed between proportions of banana genotypes within zones (S3 Table).

**Figure 1.**
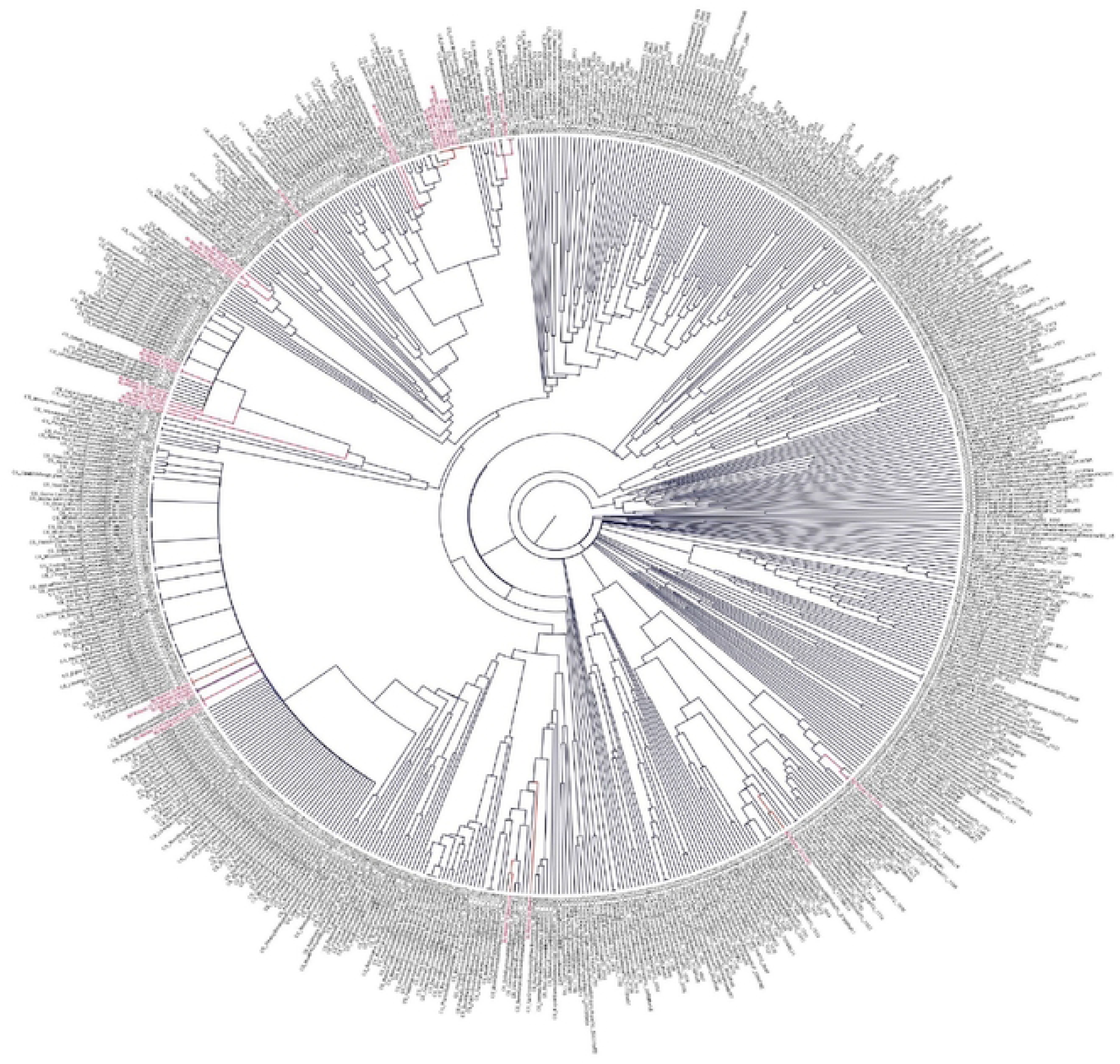
Cladogram of *Musa* germplasm diversity based on SSR markers and integrating our accessions from Malawi (labeled in red) to the banana core set collection (labeled in black) (Christelova et al. (46]).

**Table 1:** Repartition of banana genotypes (AA, AAA, AAB, AB, ABB and unknown) among the samples and prevalence of the three detected viruses (BBTV, BanMMV and BSV) for each genotype.

| Banana<br>genotype | Number<br>of<br>samples | Percentage (number) of<br>positive samples per genotype |  |  | BSV Positive to at least one of the following species: BSCAV, BSGFV,<br>BSLAC, BSIMV, BSMYV and BSOLV |  |  |  |  |  |  |
| --- | --- | --- | --- | --- | --- | --- | --- | --- | --- | --- | --- |
|  |  | Any<br>virus | BBTV + | BanMMV<br>+ | Positive to<br>all BSV<br>spp | BSCAV | BSGFV | BSIMV | BSLACV | BSOLV | BSMYV |
| AA | 5 | 40% (2) | 20% (1) | 0% (0) | 20% (1) | 0% (0) | 0% (0) | 0% (0) | 0% (0) | 0% (0) | 20% (1) |
| AAA | 67 | 42% (28) | 27% (18) | 13% (9) | 7% (5) | 0% (0) | 2% (0) | 0% (0) | 2% (0) | 4% (3) | 0% (0) |
| AAB | 26 | 46% (12) | 8% (2) | 19% (5) | 27% (7) | 0% (0) | 0% (0) | 0% (0) | 0% (0) | 31% (8) | 8% (2) |
| AB | 6 | 50% (3) | 0% (0) | 0% (0) | 33% (2) | 0% (0) | 0% (0) | 2% (1) | 0% (0) | 0% (0) | 2% (1) |
| ABB | 159 | 43% (68) | 4% (6) | 20% (32) | 26% (42) | 3% (4) | 8% (13) | 12% (19) | 0% (0) | 11% (18) | 6% (10) |
| Unknown | 8 | 38% (3) | 13% (1) | 25% (0) | 0% (0) | 0% (0) | 0% (0) | 0% (0) | 0% (0) | 0% (0) | 0% (0) |
| P value | * |  | 0.027 | 0.065 | 0.047 |  |  |  |  |  |  |

### Typology of banana seed system

The source of banana planting materials was investigated to understand Malawi’s banana seed system. This study identified six sources of banana mats namely: sharing among relatives, floods (suckers found in fields after flooding), government, Church/non-governmental organizations (NGO), own field (farm) and purchase (Table 2). Statistical analysis on the origin of the mats showed highly significant differences between sources (Chi-Sq=567.03; df = 5 and P =0.000). Sharing of propagules among relatives was by far the most popular (70 %, n=189/273), ranging between 61 to 77% depending on the zone, while NGO/Church was the lowest (0.7 %, n=1/273). A similar unbalance of banana mat origin was observed in each zone (S4 Table). Surprisingly, the results also unveiled the existence of floods (2.6 %; n=7/273) as a source of mat.

**Table 2:** Source of banana mats and prevalence of tested viruses (BBTV, BanMMV and BSV positive to the following species: BSCAV, BSGFV, BSLAC, BSIMV, BSMYV and BSOLV) within each source.

| Source of mat | Percentage (number) of mats per source | Percentage (number) of positive samples per source of mat |  |  |  |
| --- | --- | --- | --- | --- | --- |
|  |  | Any virus | BBTV + | BanMMV + | BSV + |
| Floods | 2.6% (7) | 57% (4) | 14% (1) | 43% (3) | 29% (2) |
| Government | 4.8% (13) | 46% (6) | 15% (2) | 8% (1) | 23% (3) |
| Own field | 10.3% (28) | 64% (18) | 4% (1) | 46% (13) | 21% (6) |
| Purchase | 11.8% (32) | 25% (8) | 13% (4) | 9% (3) | 9% (3) |
| Relative | 70.3% (192) | 42% (80) | 10% (18) | 14% (26) | 25% (48) |
| NGO/Church | 0.4% (1) | 100% (1) | 100% (1) | 0% (0) | 100% (1) |

Age of the mat was another parameter that was studied, and it revealed that 47 % (n=127/273) of the total banana mats were over 6 years old, followed by 39% of mats of 1-3 years old (n=106/273). The age of banana mats was not equilibrated within zones (P=0.04). For example, mats of 1-3 years old recorded 56 % (n=41/73) of the total mats within zone 1 (S5 Table) while in other zones mats of over 6 years old were the most prevalent: 47 % (32), 48 % (29) and 52 % (36) in zones 2, 3 and 4, respectively.

The types of banana cultivation system were investigated and showed that 16 % (n=43/273) of bananas in Malawi were under monoculture while 84 % (n=228/273) were under mixed cropping system, which were a majority in each cultivation zones (S6 Table).

### Virus detection

The results showed that 41 % (n=111/273) of the samples were infected with at least one banana virus (Table 3). Mixed infections were detected in 9 % (n=24/273) of the samples, corresponding to 14 samples with a dual viral infection, 8 with a triple viral infection and 2 with a quadruple viral infection. Quadruple infections of BSV (BSCAV; BSGFV; BSIMV and BSMYV) were detected in ABB genotype (Zanda cultivar) in Karonga (Zone 4). Zanda and Kholobowa. ABB genotypes from 5 sites (two in zone 1 and three in zone 3) tested positive to BanMMV, BBTV and one BSV species (BSIMV, BSOLV, BSGFV or BSMYV).

**Table 3:** Total number of samples per zone and percentages of samples tested positive for 3 viruses: BanMMV, BBTV and BSV. positive to at least one of the following species: BSCAV, BSGFV, BSLAC, BSIMV, BSMYV and BSOLV.

| Banana<br>cultivation<br>zones | Number<br>of<br>samples | Percentage<br>(number)<br>of the<br>positive<br>samples<br>per zone | Percentage<br>(number) of<br>positive samples<br>per zone |  | BSV Positive to at least one of the following species: BSCAV,<br>BSGFV, BSLAC, BSIMV, BSMYV and BSOLV. |  |  |  |  |  |  |
| --- | --- | --- | --- | --- | --- | --- | --- | --- | --- | --- | --- |
|  |  |  | BBTV + | BanMM<br>V + | All BSV<br>Species | BSCAV | BSGFV | BSIMV | BSLACV | BSOLV | BSMYV |
| Zone 1 | 74 | 50% (37) | 12% (9) | 27% (9) | 27% (20) | 3% (2) | 8% (6) | 12% (9) | 0% (0) | 12% (9) | 1% (1) |
| Zone 2 | 66 | 30% (20) | 9% (7) | 3% (2) | 20% (13) | 2% (1) | 0% (0) | 0% (0) | 2% (1) | 8% (5) | 6% (4) |
| Zone 3 | 64 | 33% (21) | 9% (6) | 5% (3) | 14% (9) | 0% (0) | 5% (3) | 2% (1) | 0% (0) | 2% (1) | 6% (4) |
| Zone 4 | 69 | 48% (33) | 10% (7) | 10% (7) | 30% (21) | 1% (1) | 15% (10) | 12% (8) | 0% (0) | 4% (3) | 16% (11) |
| Total | 273 | 41% (111) | 10.3% (30) | 12% (28) | 23.1% (63) |  |  |  |  |  |  |

BanMMV, BBTV and BSV were detected in all the four zones (S1a, 1b, 1c Fig) while no sample tested positive to CMV or BBrMV. Different species of BSV namely BSCAV, BSOLV, BSGFV, BSIMV and BSMYV were detected. Twenty-one percent (n=56/273) of banana leaf samples tested positive to one of the BSV species. The national prevalence of BanMMV and BBTV was 11 % (n=32/273) and 8 % (n=23/273), respectively.

Between zones, the prevalence of samples that tested positive for at least one banana virus was variable: zone 1 registered 50 % prevalence (n=37/74), followed by 48 % (n=33/69) for zone 4, 30% (n=20/66) for zone 2, and 33 % (n=21/64) for zone 3.

### Geographical spread of most commonly detected banana viruses in Malawi

BanMMV, BBTV and at least one species of BSV were detected in the districts of Chikwawa, Mulanje, Thyolo (zone 1); Phalombe, Thyolo, Zomba (zone 2); Dedza, Nkhotakota (zone 3) and Nkhatabay (zone 4). BanMMV was the only virus detected in Nsanje, Lilongwe and Salima while BSV was the only virus detected in Chitipa, Machinga and Mangochi. BBTV was detected in all zones although it was not detected in all districts within zones. The districts with BBTV presence were Chikwawa, Dedza, Karonga, Mulanje, Nkhotakota, Nkhatabay, Phalombe, Thyolo and Zomba (S2 Fig).

### Effect of source of mat on virus infection

Data analysis showed the source of mat significantly impacted (P≤0.05) the prevalence of BBTV, BanMMV, and BSV species (Table 2). Among all the infected mats, sharing of propagules as a source of mat had the highest prevalence of BBTV (16.2 %, n=18/111); BanMMV (17.6 %, n=26/111) and BSV (70 %, n=48/111) compared to other sources. But when considering prevalence of infection within the mat source then government, purchase and floods had higher (15.4 %; 14 % and 12.5 %) BBTV detection than 9.2 % in sharing of propagules.

### Effect of ages of banana mat on virus infection

The impact of the age of the mat on the prevalence of banana virus infection was also investigated. Data analysis for BanMMV, BBTV, BSV and total virus positive indexed samples were not significantly different (P≤0.05) between ages of mat in all zones (S5 Table).

### Effect of banana cropping system on virus infection

This study identified two banana cropping systems namely mono cropping and mixed cropping (S6 Table). Data analysis of whole survey data set and within zones revealed that total virus indexing results (BBTV, BanMMV and BSV) were not significantly different (P≤0.05) between cultivation systems.

### Impact of banana genotype on virus prevalence

The analysis of variance showed that BBTV and BSV prevalence were significantly different (P≤0.05) between genotypes while BanMMV prevalence was not significantly different between banana genotypes. Globally, AAA genotype had higher (27.3 % n=18/66) BBTV prevalence compared to ABB and AAB genotypes. All genotypes that have B genome registered higher prevalence of BSV viral infection compared to AAA genotype (Table 1). Genotype ABB recorded highest proportion 47 % (n=18/38) and 41 % (n=13/32) of BSV positive indexed samples within zones in comparison with other genotypes.

BBTV positive indexed results were significantly different (P≤0.05) between genotypes in zones 1, 2 and 4 (S3 Table). Significant differences (P≤0.05) of BSV prevalence were observed between banana genotypes only within zones 1 and 4.

## Diversity of banana viruses in Malawi

The genetic diversity of BBTV, BanMMV and BSV was studied at country-wide level. BBTV phylogenic tree (Fig 2) showed that all isolates in this study clustered with the Pacific Indiana Ocean (PIO) group and shared 100 % nucleotide (nt) identity to Malawi isolate (JQ820453) sequenced from a survey in 2012. The isolates for this study shared 99.10 % nt identity to Malawi’s isolate (ON 934241), Tonga (JF957634), Tanzania (MH795415) and Rwanda (JQ820459).

**Figure 2.**
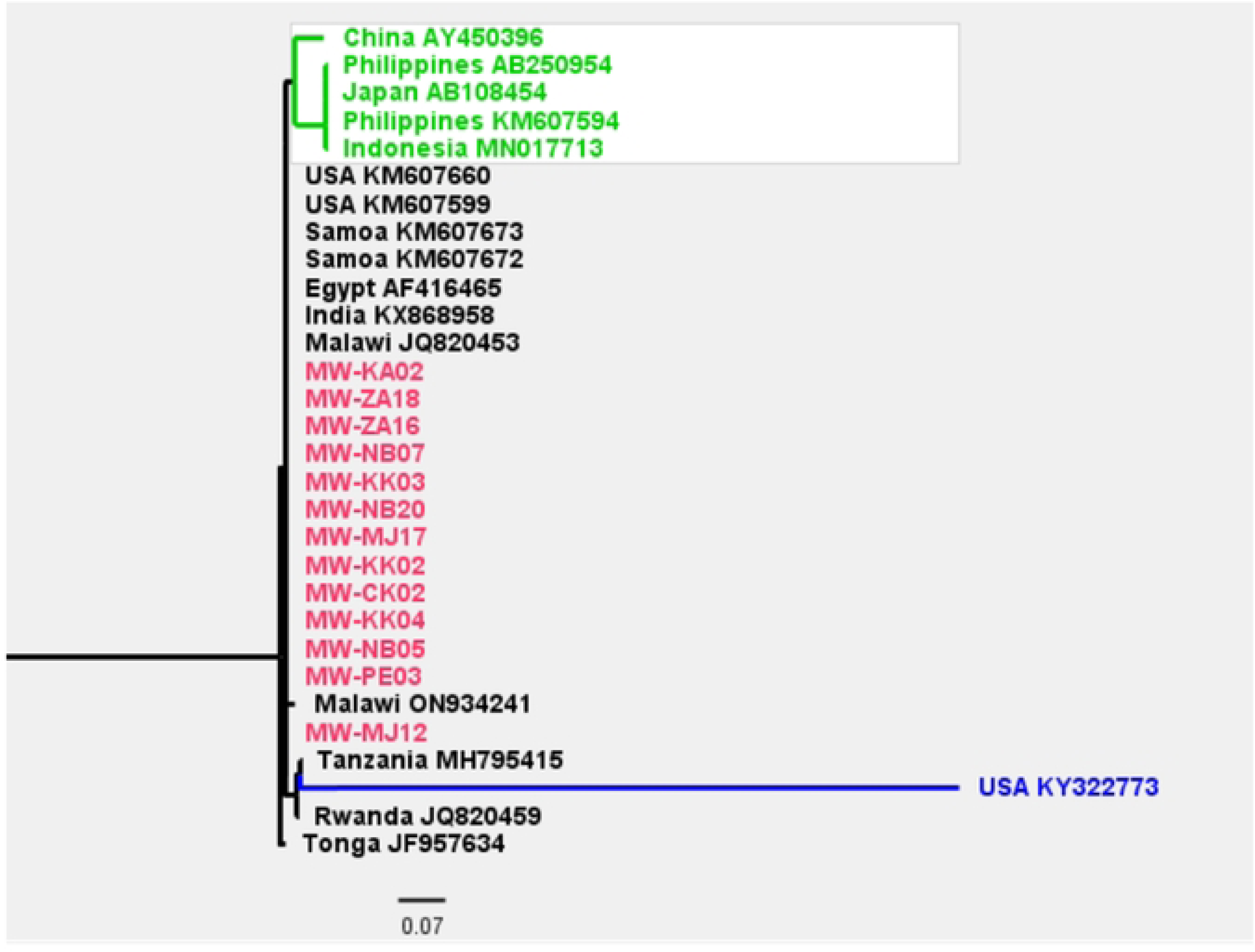
The BBTV phylogenetic tree constructed using Maximum Likelihood method and General Time Reversible model in MEGA from partial DNA-R sequences from this study (in red) and reference sequences from nt database (Genbank - NCBI, in black for Pacific Indian group, in blue for Hawaii and in green for South East Asian group).

BSV phylogenic tree showed that our isolates belonged to clade 1 (Fig 3). Isolates JM-CK5; JM-CK6 JM-CK7; JM-CP11a, JM-CP11b and KA2a shared 98.45 % identity to the Kenyan BSIMV isolate NC_015507 and 100 % nt identity to clade 1 isolate BSIMV (AJ002234). Isolates JM-CP12 and JM-KA2b had 89.92 % and 98.45 % identity to BSGFV sequence (AY493509). Two isolates, JM-DZ4 and JM-DZ8 corresponded to BSMYV and shared 92.25 % with isolate AY805074. One isolate, JM-CK8 shared 90.70 % nt identity with Cuba (FJ527427) and BSOLV (AJ002234).

**Figure 3:**
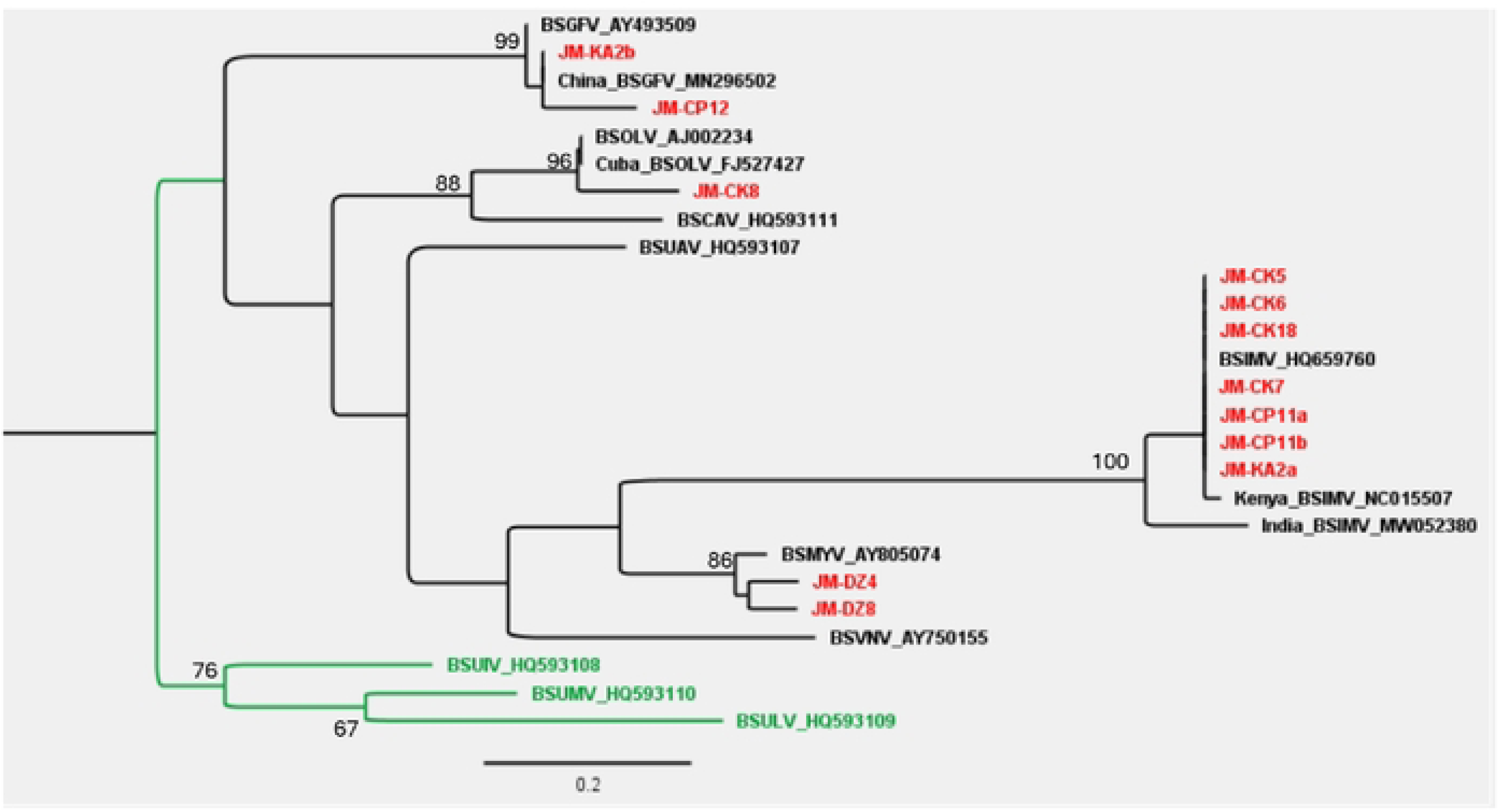
BSV phylogenetic tree constructed using Maximum Likelihood method and Jukes-Cantor model in MEGA from partial RNase H Gene sequences from this study (in red) and reference sequences from nt database (Genbank - NCBI) in black for clade 1 and green for clade 3.

BanMMV isolates JM-DZ18; JM-KK7; JM-NB25, JM-NE6 and JM-ZA10 originating from the four cultivation zones shared almost 100 % nucleotide identity on the PCR product (98.8% nt) with Australian isolate NC_002729 (Fig 4). Isolates JM-CK5; JM-LL10; JM-LL13; JM-MJ11 and JM-TO18 shared 100 % nt sequence identity to each other and with Australian isolate NC_002729. Isolate JM-MJ21 clustered with and shared 92.59 % nt to Malaysia isolate FJ179164. The pairwise identity revealed that isolates JM-PE13 and JM-NB9 shared 91.36 and 93.83 % nt sequence identity with Papua New Guinea isolates: MT872725.

**Figure 4.**
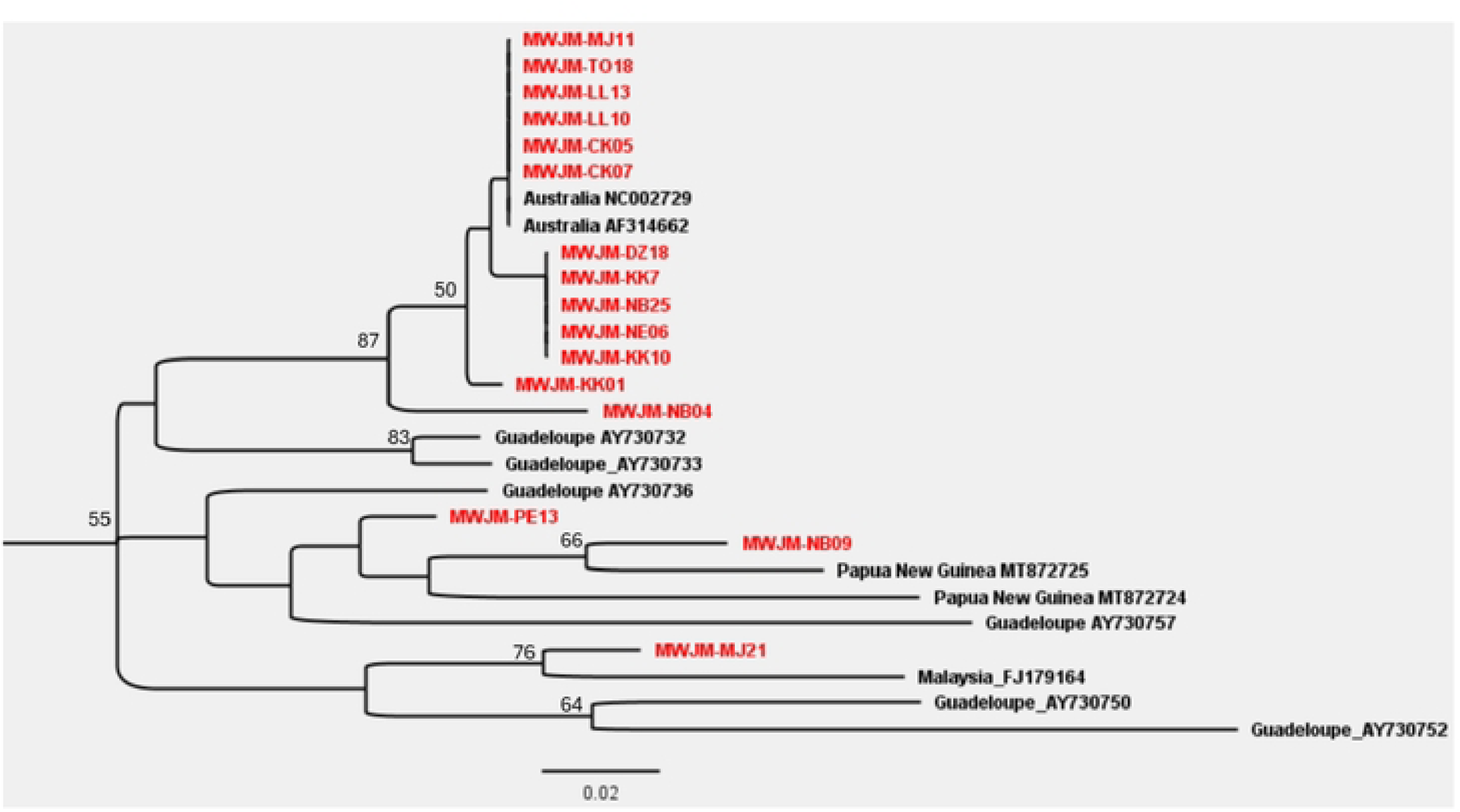
BanMMV maximum Likelihood method phylogenetic tree constructed using Jukes-Cantor model in MEGA from partial Coat protein (CP) from this study (in red) and reference sequences from nt database (Genbank - NCBI) in black.

## Discussion

Plant disease surveillance is of great importance because disease epidemics are intertwined with food security for both human and animal health [49]. For this reason, survey of banana viruses, their spread and prevalence across banana landraces within Malawi was very critical to map disease hotspot areas. This survey was also aimed at detecting pests and diseases which result in creating awareness of threats to agriculture from the entry and spread of regulated pests and pathogens [50].

In this study, the proportion of banana plants infected by at least one virus was 41%. The detected viruses were BanMMV (12 %), BBTV (10 %) and BSV spp. (23 %). Specifically, the prevalence of BSV spp. were: BSOLV (8 %); BSMIV (8 %); BSGFV (5 %); BSMyV; BSCaV (2 %) and 0.4% for BSLacV. CMV was not detected and this might be due to its low prevalence or absence on banana in Malawi. Banana bract mosaic virus (BBrMV) was not detected in this study, thus further confirming the absence of this disease in Africa compared to Asia, North America and Latin America [51]. Malawi needs to be conducting MCV and BBrMV detection surveillances to avoid epidemic emanating from the two viruses in case of their introduction. The BBTV field survey in South Africa revealed prevalence of 20 % [52]. The prevalence of BBTV from this study was 10 %, similar to that from Togo [58] but lower compared to the previous survey by Kumar *et al*. [39] which reported a BBTV prevalence of 46%. During this study six BSV species were detected, which is higher to those reported from Guadaloupe (BSOLV and BSGFV) [24] and Burkina Faso (BSLOV, BSGFV and BSIMV) [54].

Multiple infections were detected in 9 % (24/273) of the samples. Previous studies reported already the co-infection of BSV species with other banana viruses [55, 56]. Multiple infections of viruses can be important for banana virus disease management as well as sanitization of the infected plants. This implies that different techniques of sanitization of infecting viruses and also management measures need to be applied in the laboratory and at the farm level, respectively.

Importantly, the virus detection on banana samples accompanied by an extensive interview of the grower and by a characterization of the banana genotype of the plant. These data are essential for interpreting the virus incidence results.

Cultivation zones 1 and 4 had higher BBTV prevalence because of its long existence in those zones, as evidenced by Kumar *et al* [39], who detected BBTV in the mentioned zones more than a decade ago. In addition, there is a resistance of some farmers to remove the infected plants in said zones [10]. In 2018, BBTD was detected in other parts of Karonga except from its northern part which directly shares boundaries with Tanzania [11]. However, in this survey BBTV was detected in this area, meaning that BBTD is spreading to previously non-endemic areas of Malawi. In contrast to the study by Kumar *et al.* [39] that detected BBTV in samples from Lilongwe and Salima (zone 3 districts), our samples from these two districts indexed negative to BBTV. The absence of detection could be due a low prevalence or an absence of BBTV thanks to the BBTV eradication campaign by the Ministry of Agriculture and Malawi Mangoes Ltd [10, 15]. In addition, our BBTV indexing results unveiled that AAA genotype is more susceptible to BBTV than the B genomes. This study recorded the lowest proportion of BBTV positive samples within the ABB genotype despite being the dominant genome. Our findings correlated well with the findings of Ngatat *et al* [57] and Niyongere *et al*. [58] who found low infection in bananas with B genome, although no germplasm is immune to BBTV [14, 59]. In 2022, Ngatat *et al.* [57] reported that banana genotypes containing two copies of genome B have a higher tolerance to BBTV. For example, positively tested Zanda (ABB) and Kholobowa (ABB) cultivars were asymptomatic for BBTV, which can complicate eradication campaigns. The analysis of BBTV genetic diversity showed all the 13 isolates from this study clustered within the Pacific-Indian Oceans group, in agreement with previous report on BBTV isolates from Malawi and other African countries [39].

For BanMMV, its transmission is thought to occur only through propagule as no vector has been identified [60], hence its spread has most probably been facilitated by sharing infected but asymptomatic materials. The phylogenic tree, built on the partial genome sequence of BanMMV coat protein isolates, showed three sub-branches of BanMMV. Our BanMMV isolates clustered with Australia (AF314662 and NC002729), Malaysia (AY730760) and Papua New Guinea (MT872725) isolates. There was no geographical clustering of the BanMMV isolates from Malawi.

For the BSV species, our study also showed that genotypes that have B genome (AAB, AB and ABB) recorded higher BSV infection (26 % to 33 %) in all cultivation zones compared to A genomes (7 %). Our results concurred with that of Umber et al. [25], that found higher prevalence of BSV in AAB than in AAA genotypes on field and of De Clerck et al. [41] during indexing in greenhouse. This higher proportion could be due to the activation of BSV genomes (eBSOLV, eBSIMV and eBSGFV) integrated in the *Musa* B genome [21] and not the consequence of a horizontal transmission in field. These three BSV species were detected in this study together with BSCAV and BSMyV. Most samples collected in Karonga and Chitipa districts (Zone 4) indexed positive to BSV and this could be due to the very high prevalence of plantains and cooking banana (AAB and ABB genotypes) in the two districts. The banana germplasm with only A genomes were infected by BSGFV and BSOLV. BSV phylogenetic tree, based on ribonuclease H partial sequences of BSIMV, BSGFV, BSIMV and BSMYV species, showed that our isolates belonged to clade 1. The BSV genetic diversity showed that Malawi’s BSIMV isolates form a cluster with Kenyan isolate NC_015507 and had 98.17 % nt identity. The genetic diversity also revealed that our BSCAV isolates clustered with Kenyan isolate HQ593111 (100 % nt identity). The close similarities of BSIMV between Malawi and Kenya could have arisen due to sharing of propagules which is a practice in Africa [61, 62] and could be a key in the dissemination of pathogens across and within countries.

Interesting information could be gathered on the genetic diversity of *Musa* germplasm in Malawi. First, this survey revealed that Malawi Musa germplasms include five of the 10 groups defined previously [63]: AA, AB, AAA, AAB and ABB groups, with AAA, AAB and ABB being the most common. Some germplasms were unique to Malawi compared to the germplasm in the international Musa gene bank of CIAT-Bioversity Alliance. Some germplasms with different names presented a genotyping profile 100 % similar to each other, probably due to the existence of different names dependent on the site of collection or modification of the name throughout seed transmission between farmers. The banana genotype ABB was recorded in high proportion in all banana cultivation zones and this could be due to the effects of BBTD that decimated Cavendish, especially in cultivation zone 1, making those varieties carrying the B genome dominant [11, 57]. High proportion of AAA was recorded in zone 1 and the results of this study concur with Kumar *et al*., [39] who found Cavendish cultivars (AAA) popular in that zone, for fruits but also for sucker production for the commercial market as planting material. Zanda and Kholobowa were the common banana cultivars because farmers rouged off some varieties that were severely infected by BBTV leaving out Zanda and Kholobowa. Four germplasms, namely Mabere (ABB), Munowa (AAA), Kapeni (AAA) and Ndifu (AAA), were observed to be distant from other germplasm making them valuable to fill genetic gaps within the International Genebank collection.

In addition, the farmers’ interview brought interesting information to understand plant virus epidemiology. First, seed sharing amongst relatives was the main source of mat followed by sourcing from the farmer’s own field. Our findings concur with Mulugo *et al*. [64], who reported that in the Sub Sahara region, farmers prefer the informal seed system to the extent that more than 90% of farmers in the banana farming systems in Africa rely on suckers sourced from friends, neighbours, relatives and/or their own fields to establish new banana gardens. This scenario of banana planting material sharing among relatives is also supported by other studies [61, 62, 65] which reported this source as prominent in other African countries. Our results revealed that the source of mat influences BBTV, BanMMV, and BSV infection. Our results unveiled the consequence of the absence of the official banana seed system in Malawi. Sharing of propagules among relatives had the highest proportion of BBTV and BSV indexed positive samples compared to other sources. There is low availability of virus indexed propagules in Malawi which encourages sharing and poses a risk of spreading viruses.

In addition, the mats of over 6 years old were the most frequent because most smallholder farmers do not replace the old mats with new ones as long as the mats are giving them fruits. The government’s effort to revamp banana industry in areas where BBTV wiped out banana plantation had great impact which resulted in mats of 1-3 years old to be second in proportion.

High virus incidence (56 %; n=41/73 for total mats; BBTV: 15 %; n=6/41 and BSV: 34 %; n=14/41); for samples collected from plants of age group of 1-3 years old in zone 1 could be attributed to mass sharing and distribution of infected asymptomatic planting materials. Banana viruses can remain in latent form for long time and un-indexed infected mother plants used for mass multiplication can easily pass viruses to the progenies [28].

The older the plant the more time of exposure and infected with viruses, although our study did not find any effect of the age of mat on BBTV, BSV and total virus positive indexed results. No relationship between banana cultivation systems (monoculture or mixed cropping) and the prevalence of banana viruses was observed, whatever the cultivation zone.

## Conclusion

BBTV, BanMMV and BSV are the banana viruses present in Malawi while CMV and BBrMV were not detected. This study represents the first report of BanMMV and BSV from Malawi. The virus prevalence, mainly for BBTV, causes a big concern in both management and sanitization process of infecting viruses. Malawi’s BBTV isolates belong to the Pacific Indian Ocean group only and BanMMV isolates are in three sub-branches. BSV infections are high in the B genome and all the BSV spp. present in Malawi belong to clade 1. *Musa* germplasm belongs to AA, AAA, AB, AAB and ABB groups, with some germplasms unique compared to the International Genebank collection of Biodiversity International. Overall, the group ABB was the dominant one in the studied samples. Banana propagule sharing is a main source of banana planting materials in Malawi. This study found that this source promotes the spread of virus diseases, thus raising concerns for the sanitation and management of viruses and viral diseases in Malawi. There is a need of setting up good practices in the banana seed industry and policies that promote farmers’ access to virus-tested planting materials which will consequently prevent future virus epidemics. Most of Malawi’s banana mat are too old and mixed cultivation is common. Old mats need to be replaced to rejuvenate productivity. These results will assist in banana virus disease management in Malawi.

## Acknowledgments

We would like to extend our special acknowledgments to Ms. Monica Jimson for assistance rendered during banana virus survey. We acknowledge also Angelo Locicero, Igoh Lattenist and Vanessa Derycker for the technical molecular work assistance.

